# Dendritic cells accelerate CAR T cells in irradiated tumors through chimeric synapses

**DOI:** 10.1101/2024.11.19.624339

**Authors:** Sophia Navarre, Maki Ishibashi, Achuth Nair, Ivan Reyes-Torres, Meriem Belabed, Laszlo Halasz, Matthew D. Park, Raphaël Mattiuz, Merouane Ounadjela, Gertrude Gunset, Jorge Mansilla-Soto, Judith Feucht, Annalisa Cabriolu, Jessica Le Berichel, Justin Eyquem, Brian Brown, Miriam Merad, Michel Sadelain, Jalal Ahmed

## Abstract

The persistence of adoptively transferred T cells is vital for anti-tumor efficacy. Chimeric antigen receptor (CAR) T cells can persist indefinitely when delivered to patients with B cell cancers and can confer long-term remission. For patients with solid tumors, however, sustaining CAR T cell activity remains a major challenge. This has been attributed in part to the immune microenvironment within solid tumors, though the contribution of specific immune subsets to resistance to CAR T cells is not clear. Here we resolve how the immunology of irradiated tumors dramatically enhances persistence and efficacy of CAR T cells targeted to advanced lung metastases in a syngeneic mouse model. Remarkably, CAR T cell persistence depended critically on dendritic cells (DC) that underwent trogocytic “antigen-dressing” of tumor target antigens and stimulated CAR T cells through the chimeric receptor. Furthermore, tumor irradiation increased antigen-dressing onto DCs. In the absence of functional DCs, CAR T cell activity in irradiated tumor was short-lived and tumors relapsed. These findings establish a critical mechanism through which DCs maintain the CAR T cell pool in irradiated tumors, thus supporting translation of this approach to advance CAR T cell therapy for solid tumors.

## Main

Recombination of T cell signaling motifs with antigen binding receptors has been used to direct the specificity of T cells towards tumors^1^, has achieved durable remissions for patients with CD19^+^ hematologic malignancies ^2,3^, and ignited efforts to apply these Chimeric Antigen Receptor (CAR) technologies to other advanced cancers where treatment options remain limited^4,5^. In stark contrast to CAR T cells targeted to B cell malignancies, CAR T cell therapy directed to solid tumors have diminished expansion and do not persist. In a clinical trial of CAR T cells directed to Mesothelin-expressing solid tumors, for example, increasing doses of CAR T cells did not confer radiographic responses and the CAR transgene was undetectable in autopsy specimens^6^. To address this critical challenge, there has been a rapid evolution of CAR T cell technologies^7^ that venture to optimize the T cell product. To complement these efforts, we aimed to develop a better understanding of how the immunology of the tumor microenvironment (TME) dictates CAR T cell function in solid tumors.

Crucially, the chimeric receptor can provide a means for T cells to derive potent costimulatory signals from antigen presenting cells (APC) that express the CAR target, for example CD19 on B cells. DCs, the most potent APC at driving T cell responses, have successfully been engaged to expand CAR T cells. One approach enlists the endogenous TCR of virus-specific T cells (CAR-VSTs) to recognize viral peptides presented on MHC by dendritic cells^8,9^. Alternatively, RNA-LPX and Amph-ligand technologies that mark DCs with target antigens can expand CAR T cells through the chimeric receptor^10,11^ – a promising approach being tested in a phase 1 clinical trial (NCT04503278)^12^. In the absence of vaccination-based approaches, the limited persistence of solid tumor-directed CAR T cells suggested that APC-derived costimulatory signals are inaccessible in this setting, and no role for DCs was apparent. However, we observed that syngeneic 28σ CAR T cells did persist when directed to irradiated tumors, and aimed to resolve how the immunology of CAR T cells differs in irradiated tumors.

By targeting a cross-species homologous antigen (human CD19) not expressed by the host with a murine scFv (SJ25C1 mouse IgG1κ^13^) that does not bind endogenous host antigens, we intended to isolate the “On-target” activity of CAR T cells to target^+^ tumor cells. However, we were surprised to find a previously unappreciated phenomenon in which “antigen-dressing” of tumor antigens onto DCs enabled these potent APCs to expand T cells through the chimeric receptor. We quantify the impact of tumor irradiation on antigen-dressing by DCs, and provide a rationale to explore tumor irradiation as a means to advance CAR T cell therapy for patients with solid tumors.

## Results

To model the activities of immune subsets that CAR T cells would encounter in the tumor microenvironments of human patients, we used a syngeneic *K-ras^G12D^;p53^−/−^*(KP) lung tumor model shown previously using 10X single cell sequencing to develop myeloid infiltrates akin to those seen in human lung tumors^14,15^. Clonal KP cell lines transduced with the model antigen human *CD19* (*hCD19*) developed with identical kinetics to founder lines (Extended Data Fig. 1a-c) and was used for the remainder of the study. We started with the hypothesis that macrophage-depleting doses of irradiation would be necessary, and therefore selected 8Gy^16^, a well-tolerated dose used to palliate symptomatic tumors.

## Irradiation enhances CAR T cells persistence and efficacy against established lung tumors

We first assessed whether irradiation enhanced CAR T cell cytotoxicity (**Fig 1a**). 8Gy sensitized Target^+^ KP cells to CAR T cell cytotoxicity with no further benefit from higher doses (**Fig 1b**). Next, C57/B6 lung tumor-bearing mice were treated with either 8Gy of radiotherapy to the thorax (Extended Data Fig. 1d) or left unirradiated before lymphodepletion with cyclophosphamide and adoptive transfer of syngeneic CAR T cells^17^(**Fig 1a**). Irradiated mice showed significantly reduced tumor burden (**Fig 1c**), indicating enhanced CAR T cell cytotoxicity *in vivo*. Furthermore, CAR T cells targeted to irradiated Target^+^ tumors, but not Target^−^ tumors, significantly increased survival (**Fig 1d**), demonstrating a requirement for both tumor irradiation and CAR target recognition by T cells for effective treatment. Tracking of thoracic tumor bioluminescence (BLI) revealed that while CAR T cells alone or irradiation without CAR T cells both temporarily reduced tumor BLI, only irradiated mice showed a pronounced and sustained reduction in tumor BLI after CAR T cell treatment (**Fig 1e,g**) and improved survival (**Fig 1f**).

**Figure 1:**
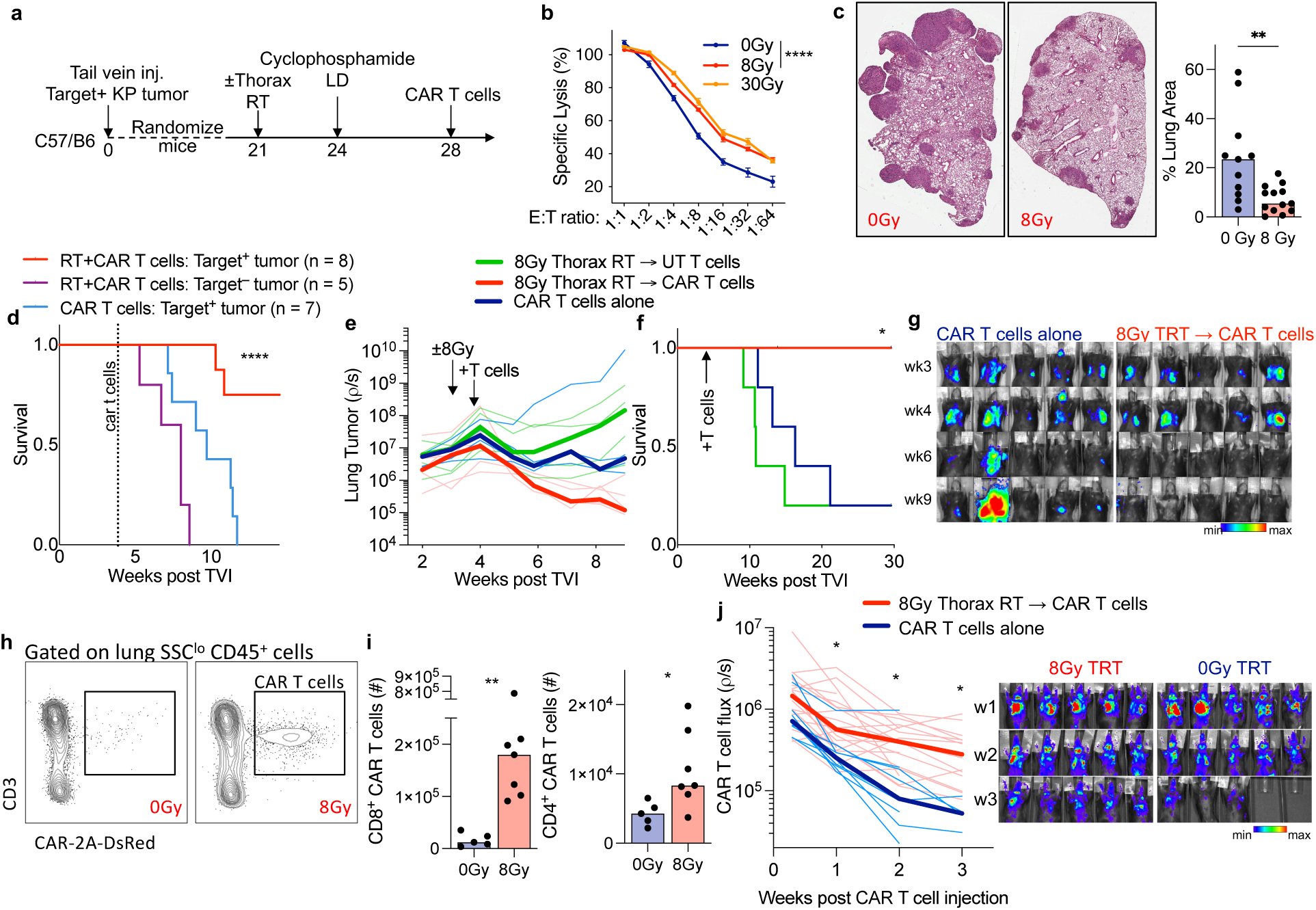
Enhanced persistence and efficacy of CAR T cells in irradiated tumors. **a, c-d,** C57/BL6 mice inoculated with 5×10^5^ target^+^ orthotopic KP lung tumors via tail vein-injection were randomized three weeks later to 8Gy thoracic irradiation or no irradiation. Cohorts were then lymphodepleted and treated with syngeneic T cells transduced with a CAR-2A-DsRed retroviral construct to a dose of 4×10^6^ CAR T cells per mouse, and lungs were harvested 9-days after CAR T cell transfer. **b,** Target^+^ Luciferase^+^ KP cells were irradiated at the doses indicated before trypsinization 8 hours later for coculture at indicated effector-to-target (E:T) ratios with CAR T cells. Mean± SEM, 12 replicates per dilution; Two-way ANOVA **** p < 0.0001. **c,** Representative hematoxylin and eosin (H&E) images of tumor-bearing lungs of mice at day 9 after CAR T cell therapy. Quantification of the tumor area as a percent of the total area of the lung cross-section (right) is shown. Each dot represents a mouse, 5-7 mice per group. Data shown compiled from two separate experiments; **p<0.01. **d**, mice were inoculated via the tail-vein with either Target^+^ or Target^−^ KP cells at a dose of 5×10^5^ tumor cells per mouse. Three weeks later, mice were irradiated or not as indicated, and then lymphodepleted and treated with 2.5×10^6^ CAR T cells per mouse. **e-g,** Thoracic bioluminescence kinetics (**e,g**) and survival (**f**) of mice injected with 5×10^5^ CBR-Luciferase^+^ target^+^ KP cells (**e-g)**. Mice were randomly allocated to treatment groups based on thoracic tumor luminescence at two weeks following tail-vein injection of tumor cells as in **1a**, and then lymphodepleted and treated with non-transduced or CAR-transduced syngeneic mouse T cells. Faint lines track tumor BLI from individual mice, median values shown in bold. **h-i,** Representative flow cytometry plots and numbers of DsRed^+^ CAR T cells in the lungs of KP19-tumor bearing mice harvested 9 days after adoptive transfer of CAR T cells. Mice inoculated with 5×10^5^ target^+^ KP lung tumors were treated with syngeneic T cells transduced with a CAR-2A-DsRed retroviral construct to a dose of 4×10^6^ CAR T cells per mouse; **p<0.01. **j,** Thoracic bioluminescence kinetics of Luciferase and CAR double-transduced syngeneic mouse T cells (2.5×10^6^ CAR T cells per mouse) injected into mice inoculated with 5×10^5^ target^+^ orthotopic KP lung tumors as in **1a**. Data combined from two independent experiments; n = 10 in 0Gy arm, n = 20 in 8Gy arm. Faint lines track T cell BLI from individual mice, median values shown in bold.

These data suggested that irradiation enhanced CAR T cell efficacy by sensitizing tumors to cytotoxicity. However, tumor BLI declined steadily (**Fig 1e-g**), suggesting sustained CAR T cell activity over several weeks. Given the ongoing challenge of maintaining CAR T cell activity in solid tumors, this was an exciting prospect that compelled us to explore whether CAR T cells were behaving differently in irradiated tumors.

Lung-tumor bearing mice were treated with syngeneic T cells transduced with CAR-2A-DsRed constructs, and numbers of DsRed^+^ CAR T cells were quantified in dissociated PBS-perfused lungs. Irradiation did not impact homing, as the numbers of CAR T cells in the lungs were similar to unirradiated mice three days after transfer (Extended Data Fig. 1e). However, nine days post-transfer, the numbers of CD8^+^ and CD4^+^ CAR T cells were 8– and 2– fold higher, respectively, in irradiated tumor-bearing lungs (**Fig 1h-i**). To track CAR T cells in individual mice, lung-tumor bearing mice were treated with syngeneic T cells double-transduced with CAR and Click-Beetle-Red (CBR) Luciferase constructs^17^. CAR T cell BLI declined rapidly in unirradiated mice. In irradiated mice, however, CAR T cell BLI diverged after the first week and remained elevated (**Fig 1j**). Evidently, the effect of irradiation was twofold: increasing cytotoxicity and enhancing CAR T cell persistence.

## Tumor irradiation induces transcriptional programs and phenotype associated with enhanced CAR T cell effector function

To identify gene signatures associated with enhanced CAR T cell activity, CAR T cells were sorted for RNA sequencing from irradiated and unirradiated lung tumor-bearing mice two-weeks post-transfer. 1585 genes were differentially expressed (DESeq2, n = 5 per group, *p* < 0.05, **Fig 2a**), of which 389 genes were enriched in CD8^+^ CAR T cells isolated from irradiated mice compared to unirradiated controls, while 1196 genes showed significantly lower expression.

**Figure 2:**
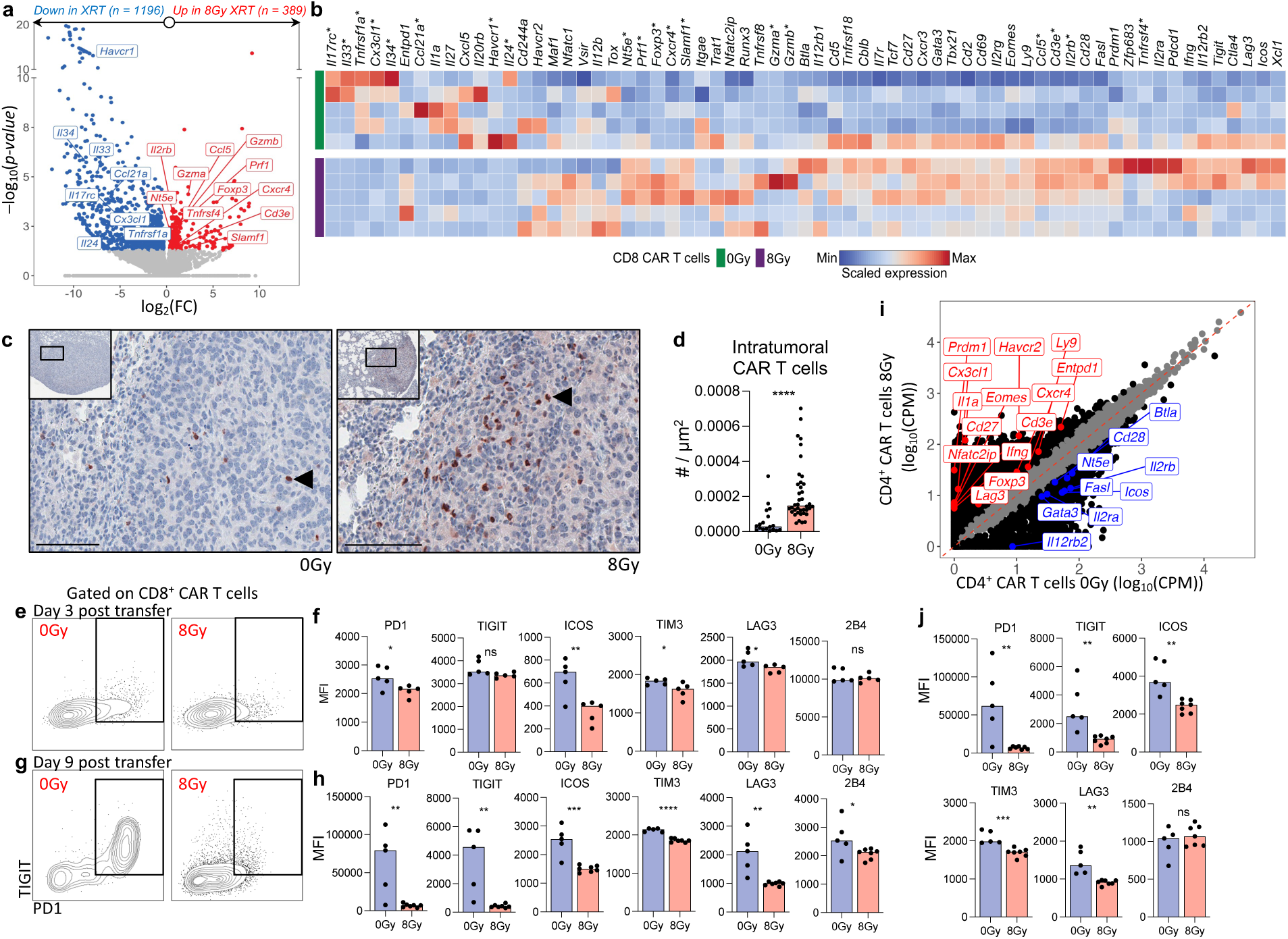
Tumor irradiation induces transcriptional programs and phenotype associated with enhanced CAR T cell effector functions. **a-b,** CD8^+^ CAR T cells were sorted from single cell suspensions of tumor-bearing lungs harvested 10-12 days after adoptive transfer (as in **1a**) and individually processed for bulk sequencing. n = 5. **a,** Volcano plot of DEGs in CD8^+^ CAR T cells from irradiated vs unirradiated mice. **b,** Heat map displayed relative amounts of transcripts of genes clustered by expression (columns) and mice (rows). Rows represent individual mice, 5 mice per group. Relative expression levels (Z-scores) of genes are shown, color-coded according to legend. * indicate significant DEG (p < 0.05). **c,** Representative paraffin sections of KP19 tumor-bearing lungs stained for DsRed expression on CAR T cells; hematoxylin counterstained. Black arrowhead indicated DsRed^+^ CAR T cell. Scale bar = 100 µm. **d,** Quantification of intratumoral DsRed^+^ CAR T cell density. Each dot represents a tumor nodule from representative lung sections from representative lung sections from 5-7 mice per group (4-10 tumors per lung section); ****p<0.0001. **e-h,** Representative cytometry plots and quantification of mean fluorescence intensity (MFI) of indicated surface markers on CD8^+^ CAR T cells 3-days (**e-f**) or 9-days (**g-h**) post adoptive transfer. **p<0.01. **i,** CD4^+^ CAR T cells were sorted from single cell suspensions of tumor-bearing lungs harvested 10-12 days after adoptive transfer (as in **1a**) and pooled (5 mice per group) before processing for bulk sequencing. Genes with < 2-fold expression between groups showed in grey, else black. **j,** Quantification of mean fluorescence intensity (MFI) of indicated surface markers on CD4^+^ CAR T cells 9-days post adoptive transfer. *p<0.05, **p<0.01

Tumor irradiation promoted expression of genes involved in ‘respiratory chain’, ‘mitochondrial protein complex’ and ‘NADH dehydrogenase complex’ Gene Ontology cell components (Extended Fig **2a**), and ‘Respiratory electron transport’ metabolic pathways, and cell cycle pathways (Extended Fig **2b**). WikiPathways ontology terms showed significant enrichment in ‘electron transport chain’, as well as enrichment of pathways involved in ‘oxidative phosphorylation’ and ‘IL-2 signaling’ (Extended Fig **2c**). These analyses suggest that CD8^+^ CAR T cells exhibit differential metabolic activity and cytokine signaling in irradiated tumors.

We compared expression of a broad set of genes implicated in T cell immunology (**Fig 2a-b**). CD8^+^ CAR T cells in irradiated tumors demonstrated elevated expression of *Il2rb*, an essential component of the IL-2 receptor complex, and transcription factor *Foxp3*, previously shown to be expressed in effector CD8^+^ T cells within human NSCLC where it was associated with higher levels of granzymes and effector cytokines^18^. Consistent with this, CD8^+^ CAR T cells in irradiated tumors were enriched in Mammalian phenotype ontology terms involved in cytolysis (Extended Fig **2d**), and key effector genes used by cytotoxic T cells to induce apoptosis in target cells (*Prf1*, *Gzma, Gzmb, Rab27a*) (**Fig 2a-b**), indicating that these cells were poised for cytotoxic function. Furthermore, the numbers of tumor-infiltrating CAR T cells were 5-fold greater in irradiated mice (**Fig 2c-d**). Therefore, irradiation increased the effector-to-target ratio within tumors, and this was associated with increased expression of genes involved in cytotoxicity and metabolism.

Although gene transcripts associated with T cell dysfunction and exhaustion had variable expression (**Fig 2a-b**), surface levels of checkpoint and coinhibitory receptors were elevated on CD8^+^ CAR T cells as early as three-days post-transfer into unirradiated mice (**Fig 2e-f**) and continued rising the following week. However, surface levels of these receptors remained low on CAR T cells in irradiated mice (**Fig 2g-h**). Similarly, CD4^+^ CAR T cells in irradiated mice did not vary in levels of checkpoint or co-inhibitory receptor transcripts (**Fig 2i**), but maintained low surface expression of these receptors (Extended Data Fig. 2e, and **Fig 2j**). These data suggest that CAR T cells in irradiated tumors downregulate co-inhibitory receptors post-transcriptionally.

## Enhanced persistence of CAR T cells in irradiated tumors requires dendritic cells

CAR T cell bioluminescence remained elevated in irradiated mice for two months after adoptive cell transfer and increased after mice were rechallenged with target^+^ KP cells (Extended Data Fig 3a). To expand and persist, T cells require strong proliferation signals from antigen-presenting cells, the most potent of which are dendritic cells (DC)^19^. Furthermore, CD8^+^ CAR T cells in irradiated tumors were significantly enriched for expression of *Ccl5*, a cytokine involved in the recruitment of dendritic cells to the tumor microenvironment^20^ (**Fig 2a-b**). We therefore hypothesized that DCs were driving the enhanced persistence of CAR T cells in irradiated tumors.

Classical dendritic cells, comprised of two distinct subsets (DC1 and DC2)^21,22^, can be distinguished from other immune lineages by expression of the transcription factor *Zbtb46*^23,24^. To time depletion of DCs, we used a knock-in mouse model in which an IRES-diphtheria toxin receptor (DTR)-mCherry cassette was knocked into the 3’-UTR of the *Zbtb46* locus^24^. The expression of *Zbtb46* on endothelial cells^23^ necessitates the generation of *Zbtb46^DTR/DTR^* bone marrow (BM) chimeras in which diphtheria toxin (DT) administration does not deplete host endothelial cells or cause lethality^24^, but depletes DCs^24^. *Zbtb46^DTR/DTR^* BM chimeras were then injected with target^+^ KP cells and randomized for CAR T cell treatment (**Fig 3a**). Remarkably, sustained depletion of DCs abolished the enhanced persistence of CAR T cells seen in irradiated mice. In contrast, CAR T cells in unirradiated mice declined with indistinguishable kinetics between DC depleted and non-depleted conditions (**Fig 3a**). These data suggest that CAR T cells do not form productive interactions with DCs in unirradiated tumors, but do so in irradiated tumors.

**Figure 3:**
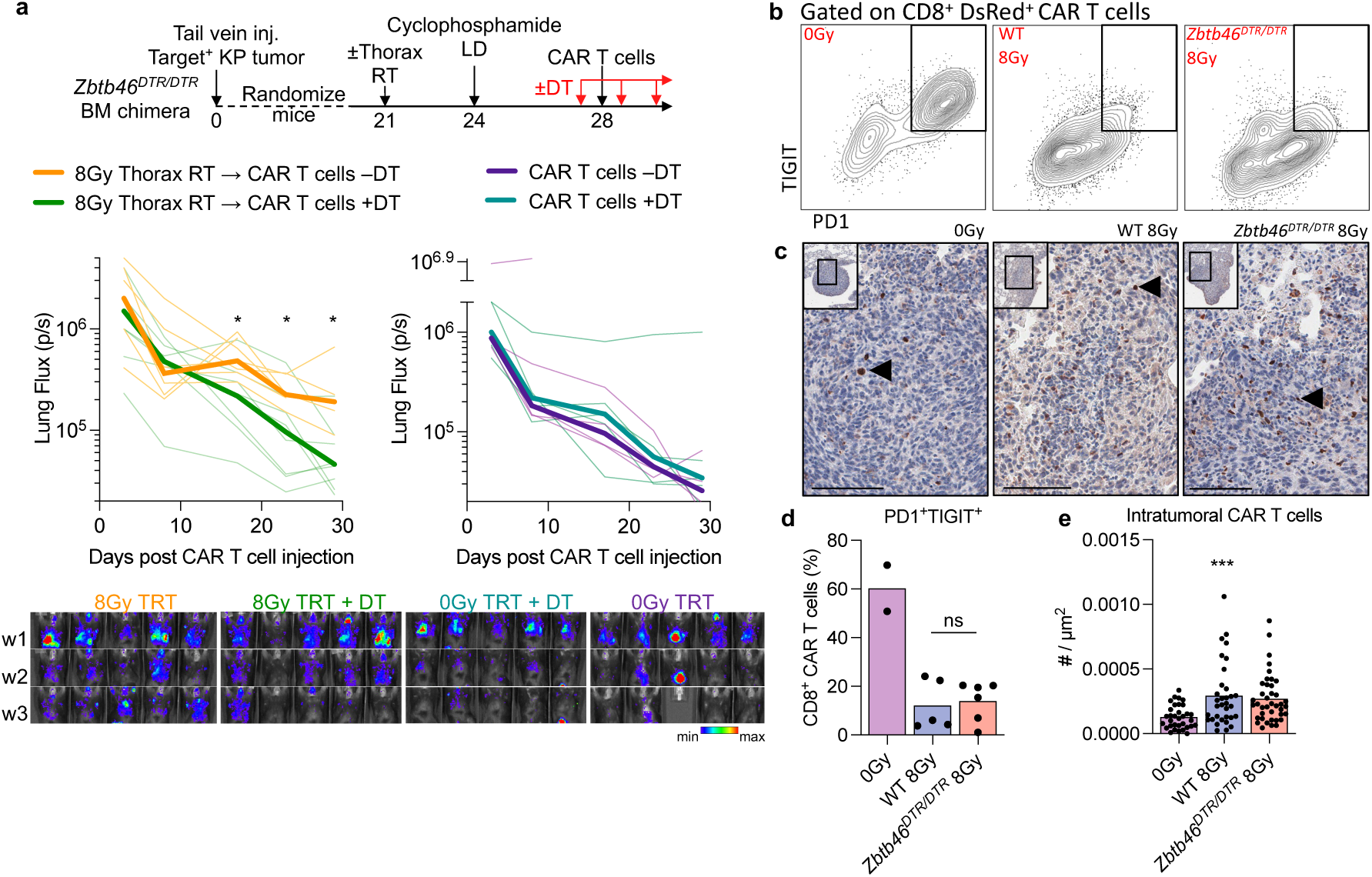
Dendritic cells are required for enhanced persistence of CAR T cells in irradiated tumors. **a,** *Zbtb46^DTR/DTR^* BM chimeras inoculated with 7×10^5^ target^+^ orthotopic KP lung tumors via tail vein-injection were randomized three weeks later to 8Gy thoracic irradiation or no irradiation. Cohorts were then lymphodepleted and treated with Luciferase and CAR double-transduced syngeneic mouse T cells to a dose of 2.5×10^6^ CAR T cells per mouse. To ensure sustained depletion of dendritic cells, Diphtheria toxin (DT) was administered starting the day before CAR T cell injection and every 2-3 days after for the remainder of the experiment in the cohorts indicated. Kinetics of thoracic CAR T cell persistence by Bioluminescence imaging were determined. Faint lines track BLI from individual mice, median values shown in bold. **b,d,** Representative flow cytometry plots and quantification (**d**) of DsRed^+^ CAR T cells in the lungs of KP19-tumor bearing mice isolated 9-days post adoptive cell transfer and stained for expression of indicated surface markers. *Zbtb46^DTR/DTR^* or WT BM chimeras were inoculated with 7×10^5^ target^+^ orthotopic KP lung tumors via tail vein-injection were then treated three weeks later to 8Gy thoracic irradiation. Cohorts were then lymphodepleted and treated with syngeneic T cells transduced with a CAR-2A-DsRed retroviral construct to a dose of 2.5×10^6^ CAR T cells per mouse. To ensure sustained depletion of dendritic cells, Diphtheria toxin (DT) was administered starting the day before CAR T cell injection and every 2-3 days after to both groups of mice. **c,** Representative paraffin sections of KP19 tumor-bearing lungs stained for DsRed expression on CAR T cells (example indicated by black arrowhead); hematoxylin counterstained. Scale bar = 100 µm. **e,** Quantification of intratumoral DsRed^+^ CAR T cell density. Each dot represents a tumor nodule from representative lung sections from representative lung sections from 5-7 mice per irradiated group (4-10 tumors per lung section); ***p<0.001 Ordinary one-way ANOVA.

Interestingly, CD8^+^ CAR T cells maintained low levels of PD1 and TIGIT in irradiated tumors whether DCs were depleted or not (**Fig 3b,d**). Furthermore, depletion of DCs did not impact CAR T cell infiltration into tumors (**Fig 3c,e**). Together, these data suggest that DCs were not driving enhanced infiltration of CAR T cells into irradiated tumors, but were required for sustaining CAR T cells over the following weeks.

## Antigen-dressed dendritic cells expand CAR T cells through chimeric synapses

Interestingly, 8Gy irradiation in the setting of lymphodepleting-chemotherapy did not drastically alter the numbers (Extended Data Fig. 3b) or transcriptional programs of DCs repopulating tumor-bearing lungs (Extended Data Fig. 3c). To assess DC and CAR T cell interactions directly, syngeneic T cells transduced with CAR-2A-mCherry-NLS^+^ constructs were added to culture wells that contained DC1 or DC2 that had been freshly sorted from KP tumor-bearing lungs, and numbers of mCherry-NLS^+^ CAR T cells were quantified on fluorescence images taken at serial time points thereafter (**Fig 4a**). DCs sorted from target^+^ KP tumor-bearing lungs had the capacity to expand CAR T cells in coculture, while DCs sorted from target^−^ KP tumor-bearing lungs did not (Extended Data Fig. 4a-b). We therefore hypothesized that target antigens on DCs were activating CAR T cells through the chimeric receptor.

**Figure 4:**
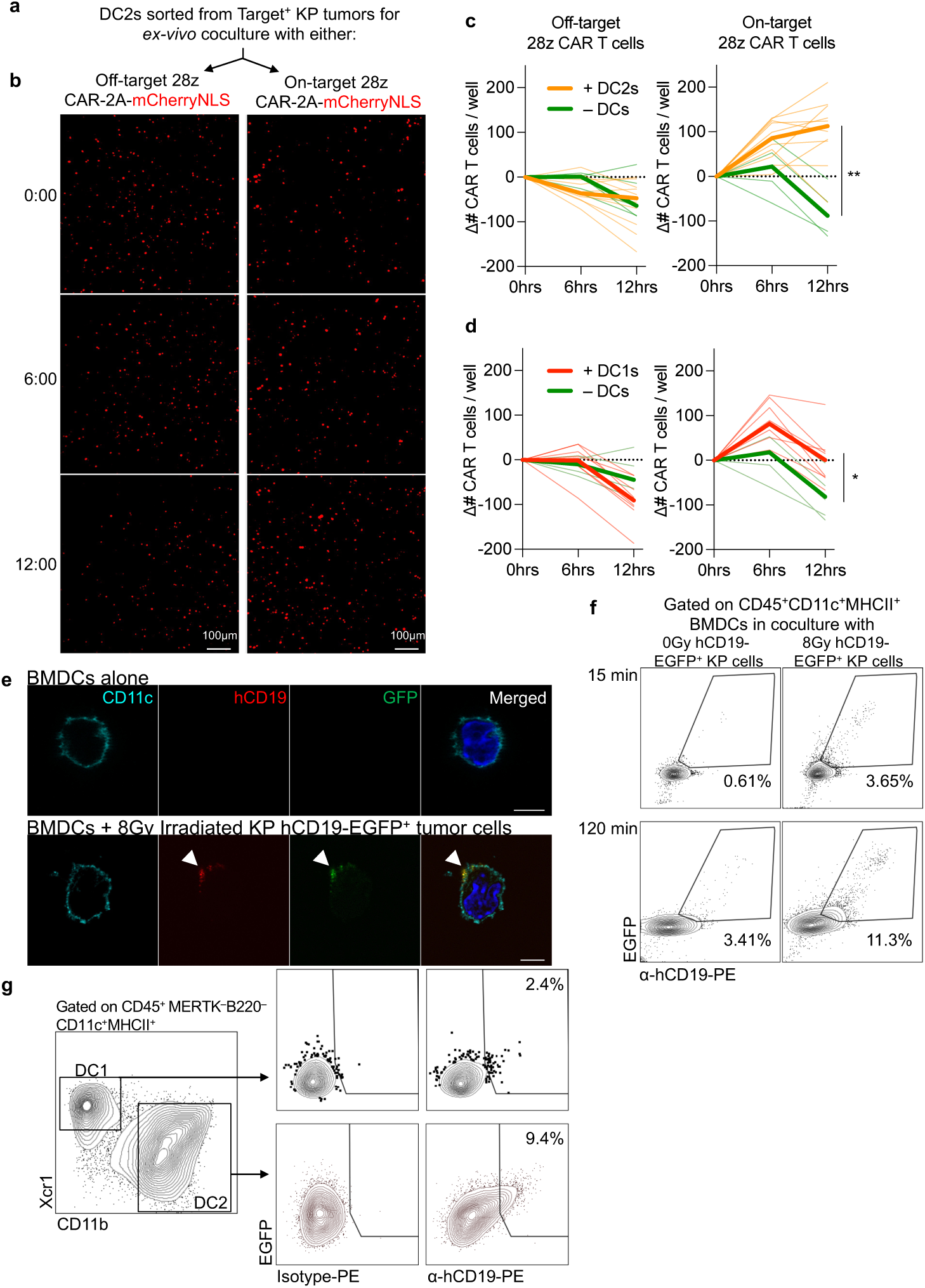
Antigen-dressed dendritic cells activate CAR T cells through chimeric synapses. **a-d**, Serial imaging and quantification of numbers of CAR-2A-mCherryNLS transduced syngeneic T cells in cocultures with DC1 (d) or DC2 (b-c) dendritic cells sorted from target^+^ KP tumor-bearing lungs, or with DC media alone. On-target (h1928z-2A-mCherryNLS) CAR T cells, and not off-target (m1928z-2A-mCherryNLS) CAR T cells, have the scFv binding domain that binds the target protein hCD19 expressed by KP19 tumors. **b**, Representative 10X widefield fluorescence images of mCherryNLS^+^ CAR T cells at the indicated time points following addition of CAR T cells to CD11b^+^Xcr1^−^ DC2s. Cells were cultured in DC media supplemented with Flt3L. **c-d**, CAR T cell numbers counted over four 10X widefield fluorescence images taken per replicate well at successive time points following the addition of “Off-target” (left) or “On-target” (right) CAR T cells to DC2 (**c**) or DC1s (**d**). Each faint line represents the change in numbers of CAR T cells per replicate well from baseline at t=0; 5 technical replicates per mouse, DCs sorted from 2 mice. Median values shown in bold. * = p<0.05, ** = p<0.01; two-way ANOVA based on general linear model with Geisser-Greenhouse correction for repeated measures. **e**, Bone marrow derived DCs (BMDC) unfed at steady state or co-cultured with 8 Gyirradiated hCD19-EGFP^+^ KP cells for 1 hour were fixed on glass coverslips and stained with anti-CD11c, anti-GFP and anti-hCD19 antibodies. Bars: 5 μm. **f**, Representative flow cytometry plots gated on CD45^+^MHCII^+^CD11c^+^ BMDCs cocultured for 15 or 120 minutes with non-enzymatically dissociated hCD19-EGFP fusion-expressing KP cells lines that were irradiated or not (24-hours prior), and stained for surface positivity of hCD19-EGFP fusion protein with α-hCD19 or isotype antibody. **g**, Representative flow cytometry plots of DC1 and DC2s isolated from KP tumor-bearing mice stained for surface positivity of hCD19-EGFP fusion protein with α-hCD19 or isotype antibody. hCD19-EGFP KP cells were injected into mice via the tail-vein and allowed to grow for 4-weeks prior to harvest.

To test the requirement of antigen recognition by the chimeric receptor, we cocultured freshly sorted DC1 and DC2s from target^+^ KP tumor-bearing lungs with syngeneic T cells transduced with either an “on-target” CAR, containing either an scFv recognition domain that binds the target antigen, or an “off-target” scFv that did not (**Fig 4a**). The addition of DCs was not sufficient to increase CAR T cell numbers in the absence of target-antigen recognition (off-target scFv) but did increase numbers of on-target CAR T cells (**Fig 4c-d**, Extended Data Fig. 4a-b).

Expansion of “on-target” CAR T cells was not due to endogenous TCR recognition of peptide-MHC presented on DCs, since “off-target” CAR T cells derived from same spleen donor have identical TCR repertoires. Therefore, these data demonstrate specificity of the chimeric receptor binding to tumor-derived target antigens on DCs.

While transfer of cell-surface proteins between immune cells has been reported in several contexts^26^, how cell surface proteins are taken up and processed by DCs, and the consequences of this for CAR T cell therapy, has not been established. To determine whether DCs could obtain surface proteins directly from tumor cells, we cocultured Bone marrow derived dendritic cells (BMDC) with non-enzymatically dissociated KP cells expressing human CD19 with EGFP fused to the intracellular domain (Extended Data Fig. 4c-d). Confocal microscopy of cocultured BMDCs visualized hCD19 and EGFP staining colocalized with CD11c on the cell membrane (**Fig 4e**). BMDCs underwent antigen-dressing from irradiated hCD19-EGFP^+^ KP cells after only 15 minutes of coculture. Tumor cell irradiation increased antigen-dressing of hCD19-EGFP onto BMDCs by 6-fold after 2 hours of coculture (**Fig 4f**). Interestingly, trypsinization of irradiated tumor cells prior to coculture abrogated antigen-dressing (Extended Data Fig 4e), suggesting that trypsin-sensitive moieties on tumor cells provide substrates for transfer of antigen to BMDCs.

Together, these data suggest that antigen-dressing can occur through trogocytosis of surface proteins, which can occur within minutes of conjugate formation between live cells, in contrast to phagocytosis and internalization that takes place over hours^27^.

To detect “antigen-dressing” *in vivo*, we injected mice with hCD19-EGFP^+^ KP cells (Extended Data Fig. 4c-d). Surface staining of live cells with anti-hCD19 antibody revealed a clear shift in EGFP-positive DCs compared to isotype-stained samples (**Fig 4g**), demonstrating that DCs undergo antigen-dressing of tumor-derived surface antigens *in vivo*.

## Sustained CAR T cell activity required for durable control of irradiated tumors

Antigen-dressing was detectable on DC1s (**Fig 4e**) that expanded CAR T cells to a lesser degree than DC2s (**Fig 4c-d**, Extended Data Fig. 4a-b). However, DC1s are critical role for cytotoxic T cell immunity^28–30^. We therefore tested whether irradiation would enhance CAR T cell efficacy in mice deficient for the transcription factor *Batf3* and lack DC1s^28^. Once again, the numbers of tumor-infiltrating DsRed^+^ CAR T cells were increased (**Fig 5a-b**) and tumor burden was reduced in irradiated *Batf3^−/−^*mice (**Fig 5c-d**). However, BLI of thoracic tumor revealed that this reduction was not sustained (**Fig 5e**). Critically, irradiation did not impact CAR T cell persistence (**Fig 5f-g**) or survival of irradiated *Batf3^−/−^* mice (**Fig 5g**). These data demonstrate that sensitization of tumors to cytotoxicity by irradiation is not sufficient, and highlight the crucial requirement for sustained activity of CAR T cells to control solid tumors.

**Figure 5:**
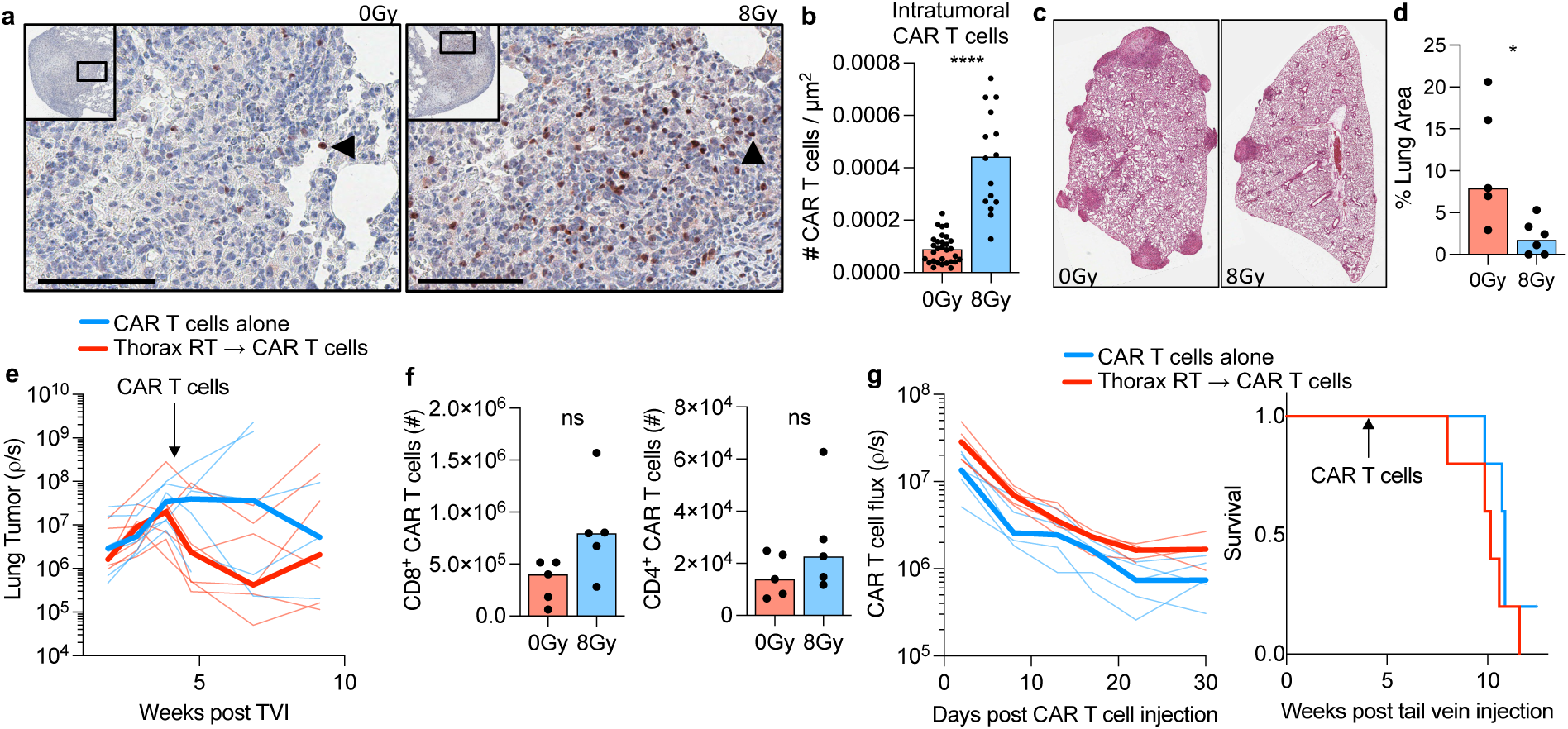
Sustained CAR T cell activity required for durable control of irradiated tumors. **a-e**, *Batf3^−/−^* mice inoculated with 5×10^5^ target^+^ orthotopic KP lung tumors via tail vein-injection were randomized three weeks later to 8Gy thoracic irradiation or no irradiation. Cohorts were then lymphodepleted and treated with syngeneic T cells transduced with a CAR-2A-DsRed retroviral construct to a dose of 4×10^6^ CAR T cells per mouse. **a,** Representative paraffin sections of tumor-bearing lungs stained for DsRed in adoptively transferred CAR T cells (selected example indicated by black arrowhead); hematoxylin counterstained. Tumor bearing lungs were harvested 9-days after CAR T cell transfer for histologic analysis. **b,** Quantification of intratumoral DsRed^+^ CAR T cell density. Each dot represents a tumor nodule from representative lung sections from 5-6 mice per group (4-10 tumors per lung section); ****p<0.0001. Scale bar = 100 µm. **c-d,** Representative H&E images (**c**) and Quantification (**d**) of tumor-bearing lungs of mice at day 9 after CAR T cell therapy. Quantification of the tumor area as a percent of the total area of the lung cross-section (right) is shown. **e,** Kinetics of Luciferase^+^ KP19 thoracic tumor bioluminescence (BLI). Mice were randomly allocated to 8Gy thoracic irradiation or no irradiation treatment groups based on thoracic tumor luminescence at two weeks following tail-vein injection of tumor cells as in **a**, and then lymphodepleted and treated with CAR-transduced WT C57/B6 T cells. Faint lines track tumor BLI from individual mice, median values shown in bold. **f,** Quantification of DsRed^+^ CAR T cells in the lungs of KP19-tumor bearing mice harvested 9 days after adoptive transfer of CAR T cells. **g-h,** Kinetics of CAR T cell persistence by thoracic BLI (**g**) and survival (**h**) in *Batf3^−/−^* mice. *Batf3^−/−^* mice inoculated with 5×10^5^ target^+^ orthotopic KP lung tumors via tail vein-injection were randomized three weeks later to thoracic irradiation or no irradiation. Cohorts were then lymphodepleted and treated with Luciferase and CAR double-transduced syngeneic mouse T cells to a dose of 2.5×10^6^ CAR T cells per mouse. n = 5. Faint lines track CAR T cell BLI from individual mice, median values shown in bold.

## α-CSF1R preconditioning does not enhance CAR T cell persistence or therapeutic efficacy

8Gy of irradiation reduced MERTK^+^ macrophages by 50% (Extended Data Fig. 5a-b). Furthermore, macrophages are the predominant myeloid population within the TME of human NSCLC^31^. To test whether targeted depletion of macrophages was sufficient to enhance CAR T cell persistence in unirradiated tumors, we administered either anti-CSF1R ^32,33^ or isotype antibody prior to adoptive transfer of syngeneic CAR T cells to mice that had established lung tumors (Extended Data Fig 5c). Surprisingly, anti-CSF1R treatment had no measurable impact on survival of mice treated with CAR T cells (Extended Data Fig 5d). Furthermore, anti-CSF1R treatment did not enhance CAR T cell cytotoxicity (Extended Data Fig 5e) or persistence (Extended Data Fig. 5f-g). However, anti-CSF1R treatment was sufficient to abrogate upregulation of PD1 and TIGIT on CAR T cells (Extended Data Fig. 5h), suggesting that thoracic radiotherapy may mitigate upregulation of checkpoint and co-inhibitory markers on CAR T cells in part through depletion of tumor associated macrophages.

## Discussion

While there are limited types of solid tumors where CAR T cell therapy have been more extensively tested including EGFRvIII^+^ or IL13Rα2^+^ Glioblastoma, GD2^+^ Sarcoma and neuroblastoma, and Mesothelin^+^ carcinomas and Mesothelioma^4^, new targets and CAR T cell constructs are continually being developed, expanding the potential application of this therapy to a broader range of solid tumors. However, durable remissions have not been achieved and CAR T cell therapy for solid tumors remains investigational. Here we implemented a clinically tractable and generalizable strategy and resolved distinct effects of tumor irradiation that contributed to enhanced CAR T cell activity.

While irradiation sensitized tumors to cytotoxicity and enhanced infiltration of CAR T cells, this was not sufficient to control tumors in *Batf3^−/−^* mice where CAR T cell activity was not sustained (**Fig 5g**), in contrast to irradiated WT recipients where CAR T cells persisted and mice achieved long-term survival (**Fig 1f**). Second, we provide the first demonstration of the requirement for antigen-dressed DCs to maintain CAR T cells in irradiated tumors. Our data demonstrates the requirement of antigen-dressed DCs not for priming of naive T cells as was shown previously with DC that were dressed with peptide-MHC from tumors ^34,35^, but for expanding CAR T cells through the chimeric receptor. In addition, we provide the first direct quantification of trogocytosis of antigens from tumors onto DCs, and demonstrate that tumor irradiation enhanced this process. Third, CAR T cells αCSF1R-treated mice maintained low surface expression of checkpoint and coinhibitory receptors, suggesting that irradiation may mitigate upregulation of these receptors in part through depletion of CSF1R^+^ myeloid cells. However, depletion of CSF1R-dependent myeloid cells was not sufficient to enhance CAR T cell cytotoxicity or persistence.

The contrast in kinetics of CAR T cells targeted to CD19^+^ hematologic cancers compared to solid tumors reflects central immunologic relationships between APCs and T cells. Indeed, labelling of target antigens onto dendritic cells expands CAR T cells several log-fold ^10,11^.

Together, these data argue that second generation CAR T cell therapies will require additional APC-derived co-signals to maintain activity against solid tumors, and suggest irradiation can further enhance efficacy in this setting.

**Extended Data Fig 1:**
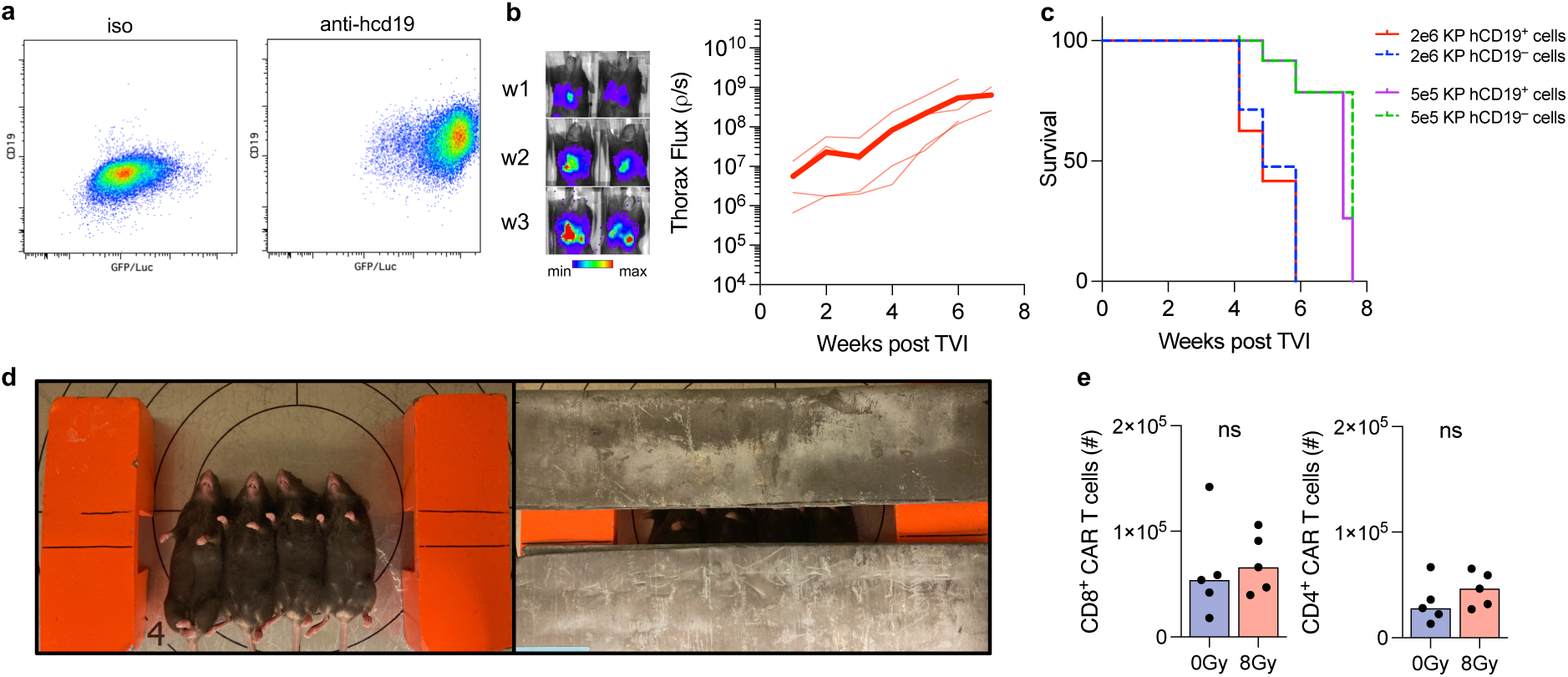
Generating a non-immunogenic lung tumor model targeted by CAR T cells. **a**, Flow cytometry of hCD19-expressing KP lung adenocarcinoma clone expressing GFP-Luciferase **b**, Bioluminescence kinetics of KP-hCD19^+^ GFP-Luciferase^+^ clonal cell line after tail-vein injection into C57/Bl6 mice, quantified in right panel **c**, Survival of mice after tail-vein injection (TVI) of KP lung adenocarcinoma cells expressing the CAR target hCD19 compared to founder KP clones that are negative for hCD19 expression (Target^−^). Two dose levels were used as indicated. Expression of hCD19 did not alter growth kinetics of KP tumor cells suggesting that this human gene did not induce an endogenous adaptive immune response. **d,** Anesthetized mice were placed supine on an aluminum shelf plate under a cone shaped irradiation field (left panel) in a RS2000 Small Animal Irradiator (Rad Source Technologies Inc.) to deliver 160kV photons at a 25mA, with a dose rate of 2.1Gy/min. Mice were treated with a half-beam block technique (right panel) eliminating divergence into the neck superiorly. Radiation was delivered to the thorax through an aperture formed between two 2mm lead sheets placed above the mice, that shielded the head and neck superior to the apex of the thorax, and tissues inferior to the xiphoid process. **e,** Quantification of numbers of CD8^+^ and CD4^+^ CAR T cells in the lungs of KP19-tumor bearing mice harvested 3 days after adoptive transfer of CAR T cells.

**Extended Data Fig 2:**
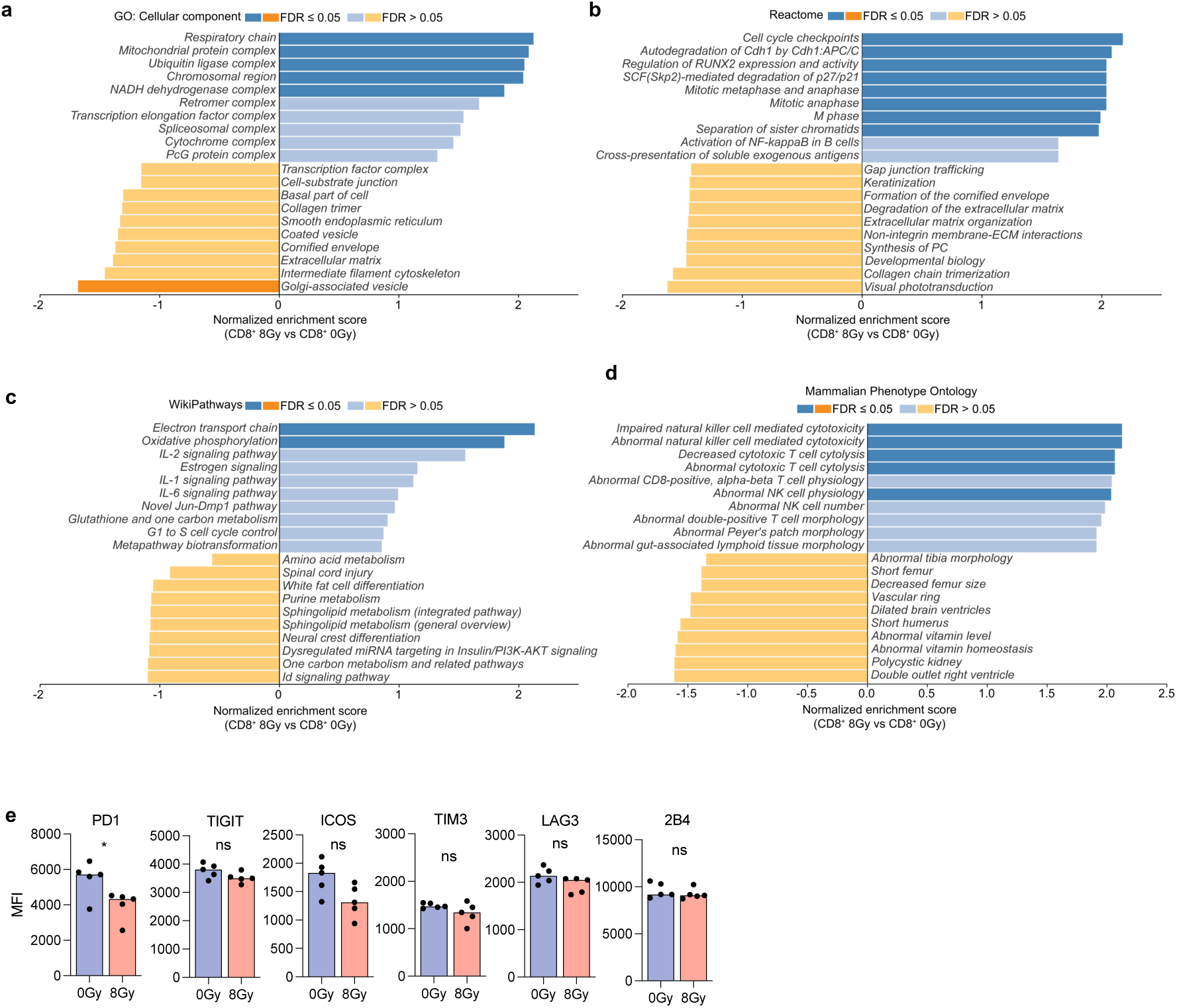
Differential transcriptional programs and phenotypes of CAR T cell incident on irradiated tumors. **a-d**, 1589 genes that were differentially expressed between CD8^+^ CAR T cells sorted from irradiated vs unirradiated mice (DESeq2, threshold of p < 0.05) were used for gene set enrichment analysis using WebGestault. **e,** Quantification of mean fluorescence intensity (MFI) of indicated surface markers on CD4^+^ CAR T cells 3-days post adoptive transfer. *p<0.05.

**Extended Data Fig 3:**
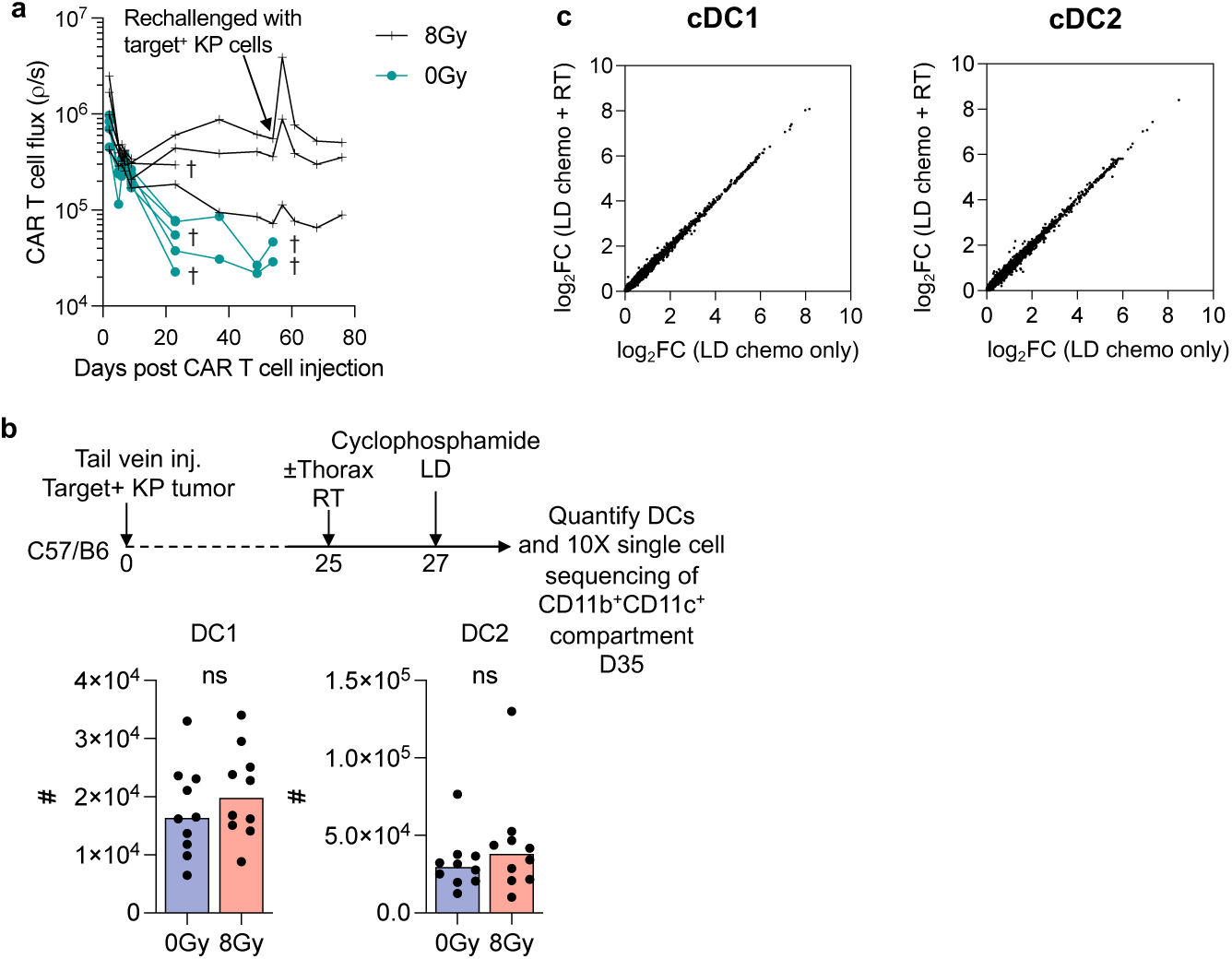
Quantification and characterization of changes in DCs in lung tumors with irradiation. **a,** Kinetics of thoracic CAR T cell persistence by Bioluminescence imaging. Mice inoculated with KP19 lung tumors were randomized to treatment with thorax radiotherapy as in **Fig 1a**, and then lymphodepleted and treated with CBR Luciferase and CAR double transduced syngeneic mouse T cells. Lines track BLI from individual mice. Arrow indicates time point when mice were rechallenged with 5×10^5^ target^+^ KP cells intravenously. **b-c,** Mice inoculated with 5×10^5^ target^+^ CBR-Luciferase^+^ orthotopic KP lung tumors via tail vein-injection were distributed based on thoracic tumor luminescence to ensure equal distribution of thoracic tumor burden to receive either 8Gy or no thoracic irradiation 25 days after tail-vein injections. They were then lymphodepleted and harvested on day 35-post tail-vein injections for quantification of dendritic cell populations (B) or for single cell sequencing of the sorted for CD45^+^CD3^−^ mCD19^−^, Ly6G^−^, CD11b ^+^ CD11c ^+^ myeloid cell compartment (C). DCs were defined as CD11b ^−^ Xcr1 ^+^ DC1s or CD11b ^+^ Xcr1 ^−^ DC2s within the following population: Live CD45^+^, Ly6G^−^, MERTK^−^, B220^−^, MHCII^+^ CD11c ^+^ **c,** Expression profiles from single cell sequencing of cDC1 and cDC2 sorted from mice treated with lymphodepleting chemotherapy alone or with 8Gy of thoracic radiotherapy. Mice were harvested 10 days after radiotherapy for single cell isolation and sorting. Plots show Log_2_FC of gene expression. Log_2_ ratios were ∼1 for all DEGs among cDC1 cells (left) and cDC2 cells (right), indicating similar transcriptome programs.

**Extended Data Fig 4:**
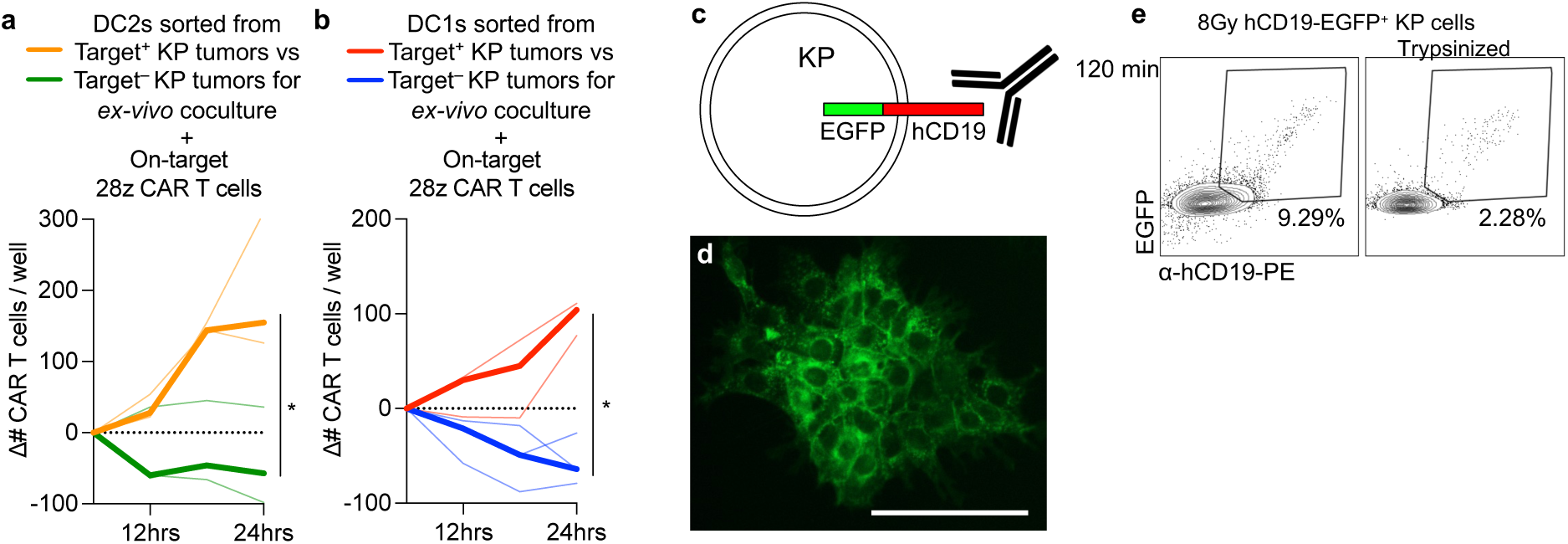
DCs sorted from target^+^ KP tumors and not target^−^ KP tumor-bearing lungs can expand CAR T cells *ex vivo*. **a-b**, Quantification of numbers of h1928z-transduced syngeneic T cells in serially-imaged cocultures with DC2 (A) or DC1 (B) dendritic cells sorted from target^+^ vs. target^−^ KP tumor-bearing lungs. On-target (h1928z-2A-mCherryNLS) CAR T cells have the SJ25C1 scFv binding domain that binds the target protein hCD19 expressed by target^+^ KP tumors, but not target^−^ KP tumor. **c,** KP tumors were transduced with lentiviral constructs expressing a human CD19-EGFP fusion molecule that allowed for detection of cell-surface vs. intracellular localization of the molecule by surface staining of cells with α-hCD19 antibody and co-expression of EGFP. **d,** fluorescence imaging of hCD19-EGFP^+^ KP clonal cell line. **e**, Representative flow cytometry plots gated on MHCII^+^CD11c^+^ bone marrow derived DCs (BMDC) cocultured for 2 hours with non-enzymatically dissociated 8Gy-irradiated hCD19-EGFP fusion-expressing KP cells. For the trypsinization condition, non-enzymatically dissociated tumor cells were pelleted and resuspended in trypsin 0.025% solution for 5 minutes then neutralized with FBS-media prior to coculture.

**Extended Data Fig 5:**
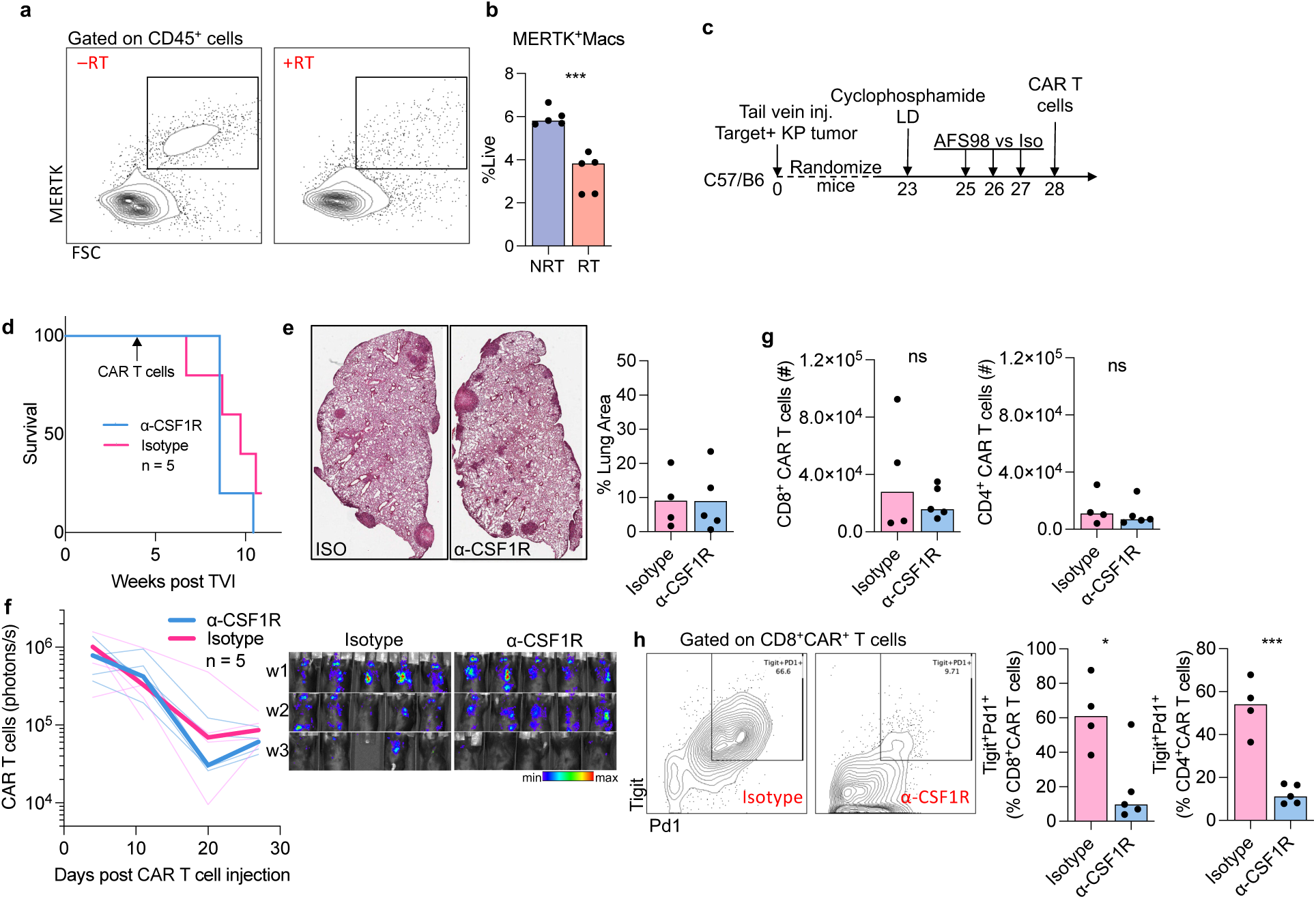
α-CSF1R preconditioning does not enhance CAR T cell persistence or therapeutic efficacy. **a-b**, Representative flow cytometry plots and quantification (B) of MERTK^+^ macrophages in KP tumor-bearing lungs. Mice inoculated with via tail vein-injection of 5×10^5^ tumor cells were randomized three weeks later to 8Gy thoracic irradiation or no irradiation. Mice were then harvested 2 weeks after radiotherapy and macrophages were quantified. **c-h,** Mice inoculated with 5×10^5^ target^+^ orthotopic KP lung tumors via tail vein-injection were lymphodepleted and randomized three weeks later to treatment with anti-CSF1R or Isotype antibody prior to CAR T cell i.v. injection. **d,** Survival of mice treated with anti-CSF1R or Isotype antibody prior to CAR T cell i.v. injection. **e,** Representative H&E images of tumor-bearing lungs of mice at day 9 after CAR T cell therapy treated with either α-CSF1R or Isotype antibody. Quantification of the tumor area as a percent of the total area of the lung cross-section (right) is shown. **f,** Kinetics of thoracic CAR T cell persistence by Bioluminescence imaging. Mice inoculated with KP19 lung tumors were lymphodepleted and randomized to treatment with anti-CSF1R or Isotype antibody as in **c**, and then treated with Luciferase and CAR double-transduced syngeneic mouse T cells. Faint lines track BLI from individual mice, median values shown in bold. **g,** Quantification of CD8^+^ and CD4^+^ DsRed^+^ CAR T cell numbers in enzymatically digested lungs of KP19-tumor bearing mice harvested 9 days after adoptive transfer of CAR T cells treated with either α-CSF1R or Isotype antibody. **h,** Representative flow cytometry plots and quantification of CD8^+^ and CD4^+^ CAR T cells isolated 9-days post transfer and stained for expression of indicated surface markers.

## Methods

### Mouse strains

Mice between 6–16 weeks of age were used following a protocol approved by the Mount Sinai Institutional Animal Care and Use Committee. Mice were maintained at specific-pathogen-free health status in individually ventilated cages at 21–22 °C and 39–50% humidity. All animal procedures were approved by the Institutional Animal Care and Use Committee (IACUC) of the Icahn School of Medicine at Mount Sinai. Mice within experiments were age and sex matched. All studies performed on mice were conducted in accordance with the IACUC at the Icahn School of Medicine at Mount Sinai.

C57BL/6J (#000664), and B6(Cg)-Zbtb46tm1(HBEGF)Mnz/J (“*Zbtb46*-DTR”, #:019506) and CD45.1 C57BL/6J-Ptprcem6Lutzy/J (#033076) mice were purchased from The Jackson Laboratory. WT C57BL/6, CD45.1, *Zbtb46*-DTR, and *Batf3^−/−^* mice were either bred at the Icahn School of Medicine at Mount Sinai or purchased from The Jackson Laboratory and housed for a minimum of 7 d before experimental use.

### Cell lines and culture conditions

KrasG12D+;p53−/− (KP) cells lines were transduced to express CAR target antigen human CD19 using Vesicular stomatitis virus glycoprotein G (VSVg) pseudotyped lentiviral supernatants derived from transfected 293 T cells transfected with SIN lentiviral transfer plasmid (pLM backbone)^36^. Stably transduced cells were single-cell sorted generate clonal cell lines prior to use in mice. KP19 clonal lines were transduced with EGFP-Click-beetle red luciferase retroviral supernatant kindly provided by Vladimir Ponomorev, to generate CBR Luciferase+ KP19 cells used for tracking tumor burden with bioluminescence. Cells were tested for Mycoplasma using PCR of DNA extracts from lysed cell pellets. Forward primer sequence: CGCCTGAGTAGTACGTTCGC. Reverse primer sequence: GCGGTGTGTACAAGACCCGA. Gel electrophoresis was used to visualize which PCR products came from myco-positive and myco-negative cells. Myco-negative cells were expanded, and batches were frozen for future use.

### Murine tumor models

To model primary lung adenocarcinoma, mice were intravenously injected via the tail vein with *Kras*^G12D/+^*p53*^−/−^ (KP) lung epithelial cells, KP cells engineered to express human CD19 (KP19), KP cells expressing both human CD19 and EGFP-Click Beetle Luciferase (CBR-Luciferase^+^ KP19), or KP cells expressing a human CD19-EGFP fusion protein (5-7 × 10^5^ cells per mouse in 200 µl phosphate-buffered saline (PBS)). KP cells were grown in complete cell culture medium (DMEM + 10% FBS + 1% P/S) and were detached for use at 70% confluence using 0.25% trypsin. The tumor cells were originally derived from KP mice^37^ generated by crossing LSL-*Kras*^G12D/+^ mice (The Jackson Laboratory) with *p53*^fl/fl^ mice (The Jackson Laboratory). Tumor quantification was performed on H&E-stained slides of formalin-fixed paraffin-embedded 4-µm lung tissue sections. Slides were scanned using an Olympus digital scanner and analyzed using the Panoramic viewer and QuPath software.

### Thoracic irradiation

Mice were deeply anesthetized using intraperitoneal injection of a mixture of ketamine (100mg/kg) and Xylazine (10mg/kg). Anesthetized mice were placed supine on an aluminum shelf plate under a cone shaped irradiation field in a RS2000 Small Animal Irradiator (Rad Source Technologies Inc.) that will deliver 160kV photons at a 25mA, with a dose rate of 2.1Gy/min.

Mice were treated with a half-beam block technique eliminating divergence into the neck superiorly. Radiation was delivered to the thorax through an aperture formed between two 2mm lead sheets suspended above the mice, that shielded the head and neck superior to the apex of the thorax, and tissues inferior to the xiphoid process. Following irradiation, mice were kept warm under a heat lamp and monitored until ambulatory.

### Lentiviral vector construction, production and transduction of cell lines

Transfer Lentiviral plasmid was generated by cloning target sequences using standard molecular biology techniques into a SIN lentiviral plasmid (pLM backbone)^36^. Vesicular stomatitis virus glycoprotein G pseudotyped lentiviral supernatants derived from transfected 293 T cells were used to transduce KP cells (see below) to generate stable KP19 cells expressing human CD19.

Transduction was performed in 6-well plates containing 4 μg ml^−1^ polybrene.

### Genetic modification of T cells

Plasmids encoding the SFGγ retroviral vector^38^ were prepared using standard molecular biology techniques as described previously^39^. Vesicular stomatitis virus glycoprotein G (VSV-G) pseudotyped retroviral supernatants derived from transduced gpg29 fibroblasts (H29) were used to construct stable retroviral-producing cell lines as described^39,40^.

SFG-h1928ζ comprises a single-chain variable fragment (scFv) specific for the human CD19 derived from the heavy (VH)– and light (VL)-chain variable regions from hybridoma cell line SJ25C1, that was cloned into an all murine CD28/CD3ζ –based CAR SFG construct. The SFG-m1928ζ construct comprises a scFv specific for the murine CD19 derived from the heavy (VH)– and light (VL)-chain variable regions from hybridoma cell line 1D3 as previously described ^41^, that was cloned into an all murine CD28/CD3ζ –based CAR SFG construct.

The anti-hCD19 scFv and anti-mCD19 scFv is preceded by a mouse CD8A leader peptide and followed by the Myc-tag sequence (EQKLISEEDL), mouse CD28 transmembrane and intracellular domain and mouse CD3z intracellular domain^40^. The SFG-h1928z-2A-DsRed, SFG-h1928z-2A-mCherryNLS, and SFG-m1928z-2A-mCherryNLS vectors were constructed by Gibson Assembly of the CAR gene without the stop codon to DsRed or mCherryNLS with the following P2A self-cleaving peptide sequence: GSGATNFSLLKQAGDVEENPGP^42^.

Spleens from euthanized 6-week-old mice were crushed and strained, RBC lysed, and T cells were enriched via negative selection using the mouse Pan T cell Isolation Kit (Miltenyi Biotec). Cells were then expanded *in vitro* by cultured in RPMI 1640 (Gibco #11875093) supplemented with FBS (10%), penicillin-streptomycin (100 U/ml), sodium pyruvate (1 mM), HEPES (10 mM), b-mercaptoethanol (Gibco #21985-023), MEM non-essential amino acids (13; Gibco #11140050) and hIL-2 (50 U/ml: Peprotech #200-02), 50 IU of recombinant human IL-2 (Peprotech #200-02), and anti-CD3/28 Dynabeads (Life Technologies) at a bead:cell ratio of 1:2. 24-h after initiating T cell activation, T cells were transduced with retroviral supernatants by centrifugation on RetroNectin-coated plates (Takara), as described previously^43^. Transduction efficiencies were determined 4 d later by flow cytometry, and CARs were either injected into tumor-bearing mice or used for in vitro experiments. For T cell imaging studies, mouse T cells were also transduced with retroviral supernatants encoding SFG-GFP-click beetle red luciferase^17,44^.

### Bioluminescence imaging and quantification

For bioluminescence tumor studies, mice were inoculated i.v. with 5 x 10^5^ CBR-luciferase^+^ KP19 tumor cells on day 0 and imaged weekly until mice reached humane endpoints.

For tracking of CBR-Luciferase^+^ CAR T cells, mice innoculated with KP19 tumor cells three weeks prior were randomized to treatment groups before undergoing lymphodepletion and intravenous injection of CBR-Luciferase^+^ CAR T cells. Cohorts were then imaged 3 days post i.v. CAR T cell injections and weekly thereafter until mice reached humane endpoints.

Mice were injected with 1.5mg of Luciferin diluted in PBS per 20g body weight and imaged immediately after. Bioluminescence imaging was done using the Xenogen IVIS Imaging System (Xenogen) with Living Image software (Xenogen) for acquisition of imaging datasets was done to guarantee equal tumor burden of mice at time of treatment. Mice were then randomized into different treatment cohorts prior to CAR T cell therapy.

### Cytotoxic Lymphocyte Assay

GFP-Luciferase expressing clonal Target^+^ KP cell lines were trypsinized and added to 96-well clear-bottom plates containing serial dilutions of CAR T cells and cocultured for 18-hour after a brief centrifugation at 1300rpm for 1min. 0.15mg in PBS Luciferin was added to cocultures immediately prior to acquisition of luminescence on plate reader. For irradiation of cell lines, culture plates at 70% confluency were irradiated on the top shelf plate under a cone shaped irradiation field in the Rad Source RS2000 irradiator.

### Adoptive Transfer of CAR T Cells

Mice were lymphodepleted 200 mg/kg cyclophosphamide (Sigma) intraperitoneally 3-4 days prior to adoptive transfer of 2.5-4 x 10^6^ CAR^+^ T cells intravenously as indicated.

### Flow cytometry and fluorescence-activated cell sorting

Mice were euthanized with CO_2_ and single-cell suspensions from perfused murine lungs were obtained upon lung tissue digestion with collagenase IV (0.25 mg ml^−1^; Sigma) at 37 °C for 30 min in agitation (80 r.p.m.) followed by passing through a 70-μm cell strainer and ACK (Ammonium-Chloride-Potassium) Lysing Buffer (Lonza) for 5 min at room temperature. For flow cytometry or FACS, cells were stained in FACS buffer (PBS supplemented with 2% bovine serum albumin (BSA) and 5 mM EDTA) with different combination of the following monoclonal antibodies (1:200 dilution) specific to: CD45 (clone 30-F11, BioLegend; cat. no. 103137); CD45.1 (clone A20, BioLegend); B220 (clone RA3-6B, BioLegend); human CD19 (clone HIB19, BioLegend); Ly6G (clone 1A8, BioLegend; cat. no. 127621); CD64 (clone X54-5/7.1, BioLegend); MerTK (clone 2B10C42, BioLegend); CD2 (clone RM2-5, BioLegend; cat. no. 100113); Siglec-F (clone E50-2440, BD Pharmingen; cat. no. 740956); MHC-I-A/I-E (clone M5/114.15.2, eBiosciences; cat. no. 12-5321-82); CD11b (clone M1/70, eBiosciences; cat. no. 45-0112-82); CD11c (clone N418, Invitrogen; cat. no. 47-0114-82); XCR1 (clone ZET, BioLegend); CD3 (17A2, BioLegend); CD8a (clone 53-6.7, BioLegend; cat. no. 558106); CD4 (clone GK1.5, eBiosciences; cat. no. 17-0041-82); CD44 (clone IM7, BioLegend); PD-1 (clone RMP1-30, BioLegend), Tigit (clone Vstm3, BioLegend). For flow cytometry, cells were analyzed in a BD LSR Fortessa analyzer (BD Biosciences) or CYTEK Aurora. For FACS, cells were prepared, stained and purified using a BD FACSAria sorter (BD Biosciences) or BD CytoFLEX SRT Cell Sorter. Flow cytometry data were acquired using FACS Diva software v.7 (BD) and the data obtained were analyzed using FlowJo (LLC).

### Imaging and quantification of CAR T cells in murine tissue sections

Mice were euthanized with CO_2_ and the left mouse lung was dissected after perfusion of the murine lungs with 10mL PBS through the right ventricle. Formalin-fixed paraffin-embedded tissue sections (4 μm) of the lungs were stained using immunohistochemical staining as previously described^45^. Briefly, slides were baked at 50 °C overnight, deparaffinized in xylene and rehydrated in decreasing concentration of ethanol (100%, 90%, 70%, 50% and dH_2_O).

Sample slides were incubated in pH 9 buffers at 95 °C for 30 min for antigen retrieval, then in 3% hydrogen peroxide for 15 min and in serum-free protein block solution (Dako) for 30 min. Primary antibody staining was performed using 1:200 dilution for 1 h at room temperature or at 4 °C overnight followed by signal amplification using associated secondary antibody conjugated to horseradish peroxidase during 30 min. Chromogenic revelation was performed using AEC (Vector). Tissue sections were counterstained with hematoxylin, mounted with a glycerol-based mounting medium and finally scanned to obtain digital images (Aperio AT2, Leica). DsRed was detected using anti-RFP (NBP1-69962 Novus Biologicals).

Tumor ROI was distinguished from lung parenchyma on corresponding paraffin sections stained with Hematoxylin and eosin (H&E). Nodules >40,000 µm^2^ were selected for quantification of intratumoral CAR T cell density. Cell segmentation was performed using StarDist extension (Github repository: https://github.com/ksugar/ stardist-sparse). Sample training images were used to train the system for accurate positive cell detection. Quantification of positively stained cells were successively performed on the ROIs using the trained algorithm and annotations were exported for downstream analysis.

Cell segmentation was performed using StarDist extension in the QuPath software (Github repository: https://github.com/ksugar/ stardist-sparse). The system was trained to accurately detect positive cells and quantification of positively stained cells were successively performed using the trained algorithm. Quantification of positively stained cells were performed using the positive cell detection algorithm in the QuPath software with default settings.

### Antibody-mediated blockade and depletion studies

To deplete CSF1R macrophages systemically prior to injection of CAR T cells, mice were intraperitoneally injected with anti-CSF1R (BioXCell, clone AFS98) or Isotype IgG2 (BioXCell, clone 2A3) over three days as previously described^46^, with the following three-day dosing schedule: 2mg day 1, 0.5mg day2, 0.5mg day 3.

### BM transplantation

BM transplantation was done as previously described^25^. Donor BM cells were harvested by canulating and flushing femurs and tibias harvested from *Zbtb46*-DTR mice and resuspended in PBS after RBS lysis. BM cell numbers were determined on Nexcelcom slides on a Cellometer Auto 2000. 10 × 10^6^ BM cells were transplanted via retro-orbital intravenous infusion.

### Diphtheria Toxin injections

Unnicked DT from Corynebacterium diphtheriae was purchased from List Biological Laboratories. *Zbtb46*-DTR bone marrow chimeras were injected i.p. with a first dose of 640 ng DT per mouse, followed by injections of 400ng per mouse every 2-3 days (Monday-Wednesday-Friday dosing schedule).

### Live cell imaging of CAR T cells and dendritic cells in *ex vivo* cocultures

C57/B6 mice were injected with 1×10^6^ target^+^ KP tumor cells via the tail-vein and whole lungs were perfused and digested to generate single cells suspensions as above. Xcr1^+^ CD11b^−^DC1 and CD11b^+^ Xcr1^−^ DC2 dendritic cell populations were sorted from Live CD45^+^Ly6G^−^B220^−^ MERTK^−^CD11c^hi^ MHCII^hi^ cells. Syngeneic T cells transduced with CAR-2A-mCherryNLS constructs as above were cocultured in 96-well flat bottom plates with DC1 or DC2 populations sorted the same day from target^+^ KP tumor-bearing lungs, or with DC media alone. DC media consisted of RPMI 1640 (Gibco #11875093) supplemented with FBS (10%), penicillin-streptomycin (100 U/ml), b-mercaptoethanol (Gibco #21985-023) and 100ng/mL final concentration of recombinant human Flt3L (carrier-free formulation, BioLegend). Plates were briefly centrifuged and then imaged at 6-hour intervals for mCherryNLS fluorescence using the BioTek Cytation 7 Cell Imaging Multimode Reader (Agilent) integrated with the BioTek BioSpa 8 Automated Incubator (Agilent). The accompanying Gen5 software (Agilent) was used to count the number of mCherryNLS-positive CAR T cells over four non-overlapping 10X widefield images captured per 96-well chamber for each timepoint.

### RNA sequencing

For bulk sequencing, CD4 and CD8 h1928z-2A-DsRed CAR T cells were sorted using the CytoFLEX SRT (Beckman Coulter) into 200 uL TRIzol LS (Invitrogen # 10296010) in DNA Lo-bind tubes (Eppendorf #022431021) pre-coated with FBS with β-mercaptoethanol 1X (Gibco #21985-023). Cells were flash frozen and submitted to Azenta Life Sciences for RNA extraction, library preparation, and bulk sequencing. Quality control, trimming, and alignment to produce a gene expression matrix from FASTQ files were done through the Human Immune Monitoring Core (HIMC) at our institute. The sequencing data was processed using the nf-core RNA-seq pipeline (https://nf-co.re/rnaseq). Briefly, we performed quality control and trimming of the raw fastq-files using FastQC and Trim Galore software, respectively. The processed fastq files were aligned to the mus musculus genome using STAR. Salmon was used to generate a gene-by-sample count matrix for further analysis. Variance stabilization transformation was performed to produce read counts, and PCA was successively performed. The count data of the transcripts were normalized using the Bioconductor package DESeq2. Differential gene expression analysis was also performed using the DESeq2 package with a threshold of *P* value less than 0.05. Genes of interest were aggregated and their expression standardized with z-scores across the samples in order to produce the heat map. We performed over-representation analysis (ORA) using the WEB-based Gene SeT AnaLysis Toolkit (WEBGestalt), version 2019.

For 10X Single Cell Sequencing, CD11b^+^CD11c^+^ double positive fraction was sorted from sorted from CD45^+^CD3^−^ mCD19^−^, Ly6G^−^ fraction of single cell suspension from tumor-bearing lungs ten days after radiotherapy was delivered to half the cohort. Mice were irradiated 3 weeks after tail-vein injection of Target^+^ KP cells. Both cohorts underwent lymphodepletion 7 days prior to sorting. Live cells were submitted to the Mount Sinai Human Immune Monitoring Core (HIMC) for single cell sequencing on the 10X platform. Libraries processed with Cell Ranger v6.1.2 and aligned to reference refdata-gex-mm10-2020-A.

### BMDC coculture

To generate mouse bone marrow-derived dendritic cells (BMDCs), bone marrow precursors were cultured in IMDM medium supplemented with 10% fetal calf serum (FCS), 20 mM glutamine, 100 U ml−1 penicillin–streptomycin, 50 μM 2-mercaptoethanol (2-ME), and supplemented with granulocyte-macrophage colony-stimulating factor (GM-CSF) at a concentration of 50 ng ml−1, derived from supernatant obtained from transfected J558 cells, for a period of 9 days, as per established protocols. Immature dendritic cells were isolated by gently retrieving semi-adherent cells from the culture dishes and added to vials containing non-enzymatically digested tumors cells(using Cellstripper™, Corning) at a ratio of 1:10.

For confocal imaging of antigen-dressed BMDCs: BMDCs were cultured on glass coverslips and fixed by incubating for 15 minutes on ice in 3.7% w/v paraformaldehyde. The cells were then quenched for 10 minutes with 50 mM NH4Cl in PBS. Subsequently, the cells were incubated for 1 hour with specific primary antibodies in PBS, washed twice, and incubated for an additional hour with fluorescently labeled secondary antibodies. Finally, the cells were mounted on slides using Prolong Gold antifade reagent with DAPI. Confocal microscopy was performed using a Zeiss LSM 780 system (Carl Zeiss) with 63x objectives. Image processing was carried out with Zeiss LSM Image Browser (Carl Zeiss) and ImageJ software.

## End notes

## Acknowledgments

We thank the following for discussions and support of the authors over the course of this study: M. Casanova, L. Troncoso, C. Park, N. Tung, P. Hamon, the Human Immune Monitoring Center at Mount Sinai: T. Dawson, D. D’Souza, S. Kim-schulze; D. Filipescu for guidance and technical expertise in the development of live cell imaging assay; G. Ioannou, for technical expertise in the development of live cell imaging assay; J. Mansilla-Soto, A. Cabriolu, J. Feucht, M. Hamieh, Z. Zhao, A. Dobrin, J. Eyquem, A. Odak, N.Jain and G. Gunset from the Sadelain lab for providing reagents, technical expertise, and discussions in the development of the CAR T cell models used in this study.

## Author contributions

J.A.K, S.N, M.I., A.N. contributed to vector construction, T cell transduction, cell culture, flow cytometry and animal studies, and compiling of the manuscript.

M.I., A.N., performed immunohistochemistry stains and imaging

S.N., M.B., M.B. performed BMDC coculture assays and confocal imaging.

M.I. performed spatial quantification analyses of tissue sections.

A.N. performed cell sorting for single cell sequencing.

I.RT., contributed to animal studies, design of flow panels for myeloid cells. M.I., M.D.P. performed single cell sequencing computational analyses.

R.M., J.MS., M.B., B.B., provided intellectual input and technical expertise. J.MS., J.F., L.C., G.G. contributed to vector design and production

M.M, M.S. contributed to conceptualization and data interpretation

J.A.K., conceived the project, supervised the studies, analyzed and interpreted data and wrote the manuscript.

## Data and materials availability

All data are available in the main text or the supplementary materials except for sequencing FASTQ files which will be uploaded to GEO.

Correspondence and requests for materials should be addressed to

